# Factors influencing knowledge and practice on helping babies breathe among Skilled Birth Attendants in rural areas in Lake Zone in Tanzania

**DOI:** 10.1101/462903

**Authors:** Cecilia B Mzurikwao, Secilia K Ng’weshemi Kapalata, Alex Ibolinga Ernest

## Abstract

**Background:** It is estimated that 1 million babies die each year due to birth asphyxia. Globally, it is approximated that 10 million babies cannot do it by themselves and need assistance. Helping babies breathe is a key component in reducing neonatal mortality due to birth asphyxia.

**Methods:** A cross-sectional design was used, A total of 330 respondents included in the study. Simple random sampling by lottery was used to select the 2 regions and health facilities. The participants were selected through convenient. Data were collected using standard semi-structured questionnaire. Chi-square and Binary logistic regression were used to analyse the data.

**Results:** Out of 330 participants, Those who working in hospital and were more likely to have adequate knowledge (AOR= 3.227, P< 0.001) and practice (AOR= 43.807, P<0.001) than those working in Health Centers; Enlored nurses were more likely to have adequate knowledge (AOR= 3.118,P<0.05) than AMO/MD;Those with 1 year and above of experience in labor ward were more likely to have adequate practice(AOR=15.418,P<0.001) than those with less than 1 year of experience in labor ward; those who attended once on neonatal resuscitation training were adequate knowledge (AOR=1.778,P<0.05) than those who had never attended. Those with Enough equipment of neonatal resuscitation had adequate practice (AOR=4.355, P<0.001) than with no enough equipment.

**Conclusion:** Regarding the findings of the current study, it was revealed that working facility, Professional/ qualification, and training was significant predictor of knowledge while working facility, experience, and equipment was significant predictor of practice. There is a need to find effective measures on how to reduce those factors which affect knowledge and practice on helping babies breathe.

## INTRODUCTION

The health of new born babies is very important during delivery in daily life, and it is estimated that more than 100 million babies are born annually worldwide(1). During the process of birth, babies can make the transition from the intrauterine life to extra uterine life. During the intrauterine life, the baby can use the placenta as an organ for gaseous exchange, and soon after birth, the fetus can use their own cardiopulmonary system independently within the minutes of survival.

It is estimated that all babies do not establish their own spontaneous breathing during birth, so helping babies breathe is needed as an intervention for neonatal resuscitation during delivery and can be done by skilled birth attendants(1).

Almost 99% of all the deaths take place in resource-poor settings (2), A major factor contributing to the high mortality is a global lack of trained providers in neonatal stabilization and/or resuscitation which is most acute in Sub-Saharan Africa with the highest neonatal mortality (2).

Helping babies breathe is an evidence-based education programme that was developed by the American – Academy of Pediatrics in consultation with the WHO and its collaborative partners to teach neonatal resuscitation technique in resource-limited areas to health care providers in developing countries(3). The goal is to have at least one skilled birth attendant at each delivery and to reduce the neonatal mortality rate due to birth asphyxia. This essential intervention is useful by providing them with stimulation to breathe, within one minute hence many babies can be saved (4).

Estimated that one million of babies die each year due to birth asphyxia (3), Birth asphyxia is the inability of a new born baby to breathe immediately after birth. Globally, it is approximated that 10 million babies cannot do it by them and need some assistance at birth(3). Therefore helping babies breathe can focus on helping them initiate breathing. It was found that 99% of babies need this intervention, almost 80-90% who need this will be sufficient (WHO, 2012) as cited by (4).

Globally, about one-quarter of all neonatal deaths are caused by birth asphyxia and is among of the leading cause of those neonatal deaths, which account 23%( Lawn et al.) as cited by (5). The neonatal resuscitation has the potential to prevent perinatal mortality caused by birth asphyxia for almost two million babies annually (Lawn et al.) as cited by(5),and it is estimated that 8.2 million of children under five die each year, 3.3 million deaths occur in the first four weeks of life and the neonatal deaths account for an increasing proportion of children’s deaths by 41 % (Lawn et al.) as cited by (5).

The main causes of these deaths are an infection (25%), birth asphyxia (24%) and complication of pre-term 25 % (6). In Tanzania, the neonatal mortality rate due to birth asphyxia accounts for 31% (7). This unacceptably high rate must be reduced if the success in achieving better child survival is to be reached. In low–income countries, nearly half of all new born do not receive skilled care during and immediately after birth(7). However, two thirds of new born deaths can be prevented if effective measures are taken(7). Neonatal resuscitation (NR) is a simple, inexpensive intervention that has been shown to reduce neonatal mortality(7).

Due to the slow decline of the Millennium Development Goal 4 which ended in 2015 to reduce the neonatal mortality rate, the Sustainable Development Goal targets 3.2, strategies aimed to reduce new born deaths at least to 12 deaths per 1000 live births by 2030 (Lawn et al) as cited by (5).

The Helping Babies Breathe programmes offer standardized training, knowledge, and skills in essential new born resuscitation to providers and have the potential to significantly reduce global neonatal mortality. Since its introduction in 2010, HBB has been taught in 77 countries to approximately 300 000 birth attendants and is currently being implemented throughout sub-Saharan Africa, Asia, and Latin America (8).

However, the data on its effectiveness at reducing neonatal mortality are interpreted differently. The falloff of skills and knowledge after training in new born resuscitation has been well documented in high-income countries(8). Nevertheless, in the low-income countries, there are few data. Furthermore, there is poor retention of knowledge and skills after training courses which represent a significant barrier to improving neonatal mortality worldwide (8).

Helping Babies Breathe is initiated soon after delivery where all new-borns are evaluated for crying. That is, if the baby doesn’t cry, it will need basic resuscitation. The intervention includes thoroughly drying the baby and stimulating it to breathe, cleaning the airway starting from the mouth then to the nose to avoid aspiration. This is followed by the bag and mask ventilation. However, few babies’ approximately one percentage will need advanced resuscitation including specific medications and cardiac resuscitation. However, this intervention has been under-utilized due to a shortage of well-trained staff and a shortage of equipment in the resource-limited area, which includes Tanzania (9).

In 2009, a simulation programme of helping babies breathe was implemented to eight hospitals in Tanzania. These included three major referral hospitals (Muhimbili National Hospital, Bugando Medical Centre and Kilimanjaro Christian Medical Centre), four regional hospitals (Amani, Burunguni, Sekou Toure and Mawenzi), and one district hospital (Haydom Lutheran Hospital). These hospitals were successful in reducing new born deaths within 24 hours as well as reducing stillbirth (9)

In April 2010, the initial one-day HBB course was held at Hydom Lutheran Hospital. Only half of the birth attendants were able to attend and no local Master Instructors were trained to facilitate the continuation of HBB re-trainings. An evaluation of data from Hydom Lutheran Hospital revealed no improvements in mortality during the first months after the initial HBB course (2).

An additional study on testing skills and knowledge among birth attendants at Hydom Lutheran Hospital showed that both knowledge and technical skills improved when rated in a simulated setting seven months after the initial one-day HBB course compared to what it was before. However, this improvement did not transfer into clinical practice (10).

Despite the neonatal resuscitation programme which was successfully initiated, birth attendants (BAs) must be assessed on their knowledge and practice to perform appropriate and adequate neonatal resuscitation in the critical first minutes after birth (11).

## METHOD AND MATERIAL

### Research Design

The analytical cross-sectional study was used with quantitative approach. This is because the analytical cross-sectional study was used to investigate the relationship between exposure and outcome of interest. The data were collected at one point in time.

### Research Setting/Area

This study was conducted in rural areas in Lake Zone in Tanzania Mainland. Lake Zone consists of 4 regions namely Kagera, Mwanza, Mara, and Geita. In particular, 2 regions namely, Mwanza and Kagera Region were used for the study. Specifically, 5 district hospitals from each Region and 10 Health Centre’s located in a rural area were studied.

The referred samples which have been studied were composed of both Government and Non-Government Hospitals.

Mwanza Region has 7 districts namely Ilemela, Magu, Nyamagana, Kwimba, Misungwi, Sengerema, and Ukerewe. Mwanza Region has a population of 2,772,509 (Tanzania National Census of 2012). The five districts in Mwanza Region involved in the study included Magu, Sengerema, Kwimba, Nyamagana, and Misungwi. Whereby in each District, 1district hospital and 2 health centers were studied(12).

The study was also conducted in Kagera Region, whereby 5 districts namely Ngara, Karagwe, Bukoba Rural, Kyerwa, and Muleba were involved. According to Tanzania Population and Housing Census (2012), the region has a total population of 2,458,023. Health services are being provided to all the regional population with about 15 hospitals available in the region distributed in all districts. Also in Kagera region,1 district hospital and 2 health centers were used in each district.(13).

### Study Population

The study population included the skilled birth attendants who were conducting deliveries in the labor ward.

### Inclusion Criteria

The inclusion criteria focused on all skilled birth attendants such as AMO, medical doctors and nurses midwives with certificate, diploma, undergraduate and master degree, who were conducting deliveries in labor ward within the time of data collection and who gave their informed consent by agreeing to participate in the study.

### Exclusion Criteria

The exclusion criteria focused on those skilled birth attendants who were sick, absent, student nurses and doctors and those who did not agree to participate during the data collection day.

### Sample Size

The sample size was estimated using the Kish Leslie’s formula for quantitative studies (14) and the prevalence of current evaluation of birth asphyxia in Tanzania 31% (7).

**The formula states that;**

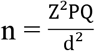

n = minimum sample size

Z = Constant, Standard Normal Deviance

(1.96 for 95% confidence level)

P =31%= prevalence of birth asphyxia (7)

Q= (1-p)

d = Acceptable Margin of error 5%

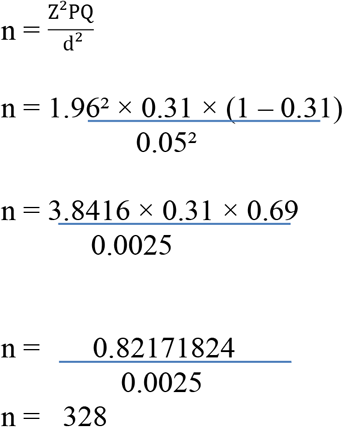

### Sampling Techniques/Method

Simple random sampling by lottery was used to select 2 regions (Mwanza and Kagera). A total of 5 district hospitals from rural Kagera and 5 of them from rural Mwanza were selected by simple randomly by lottery. Also, a total of 10 district health centres from rural Kagera and 10 of them from rural Mwanza were selected randomly and through lottery selection in all districts. The researcher prepared written small pieces of paper and then the research assistant picked them. (That is, 2regions, 5 district hospitals in each region and 2 health centres from each district in both regions).

The participants were selected through convenient sampling from those district hospitals and health centres in both Mwanza and Kagera Region, the number of participants in each region was estimated according to the number of health facilities, and staff working in the labour ward, where 17 participants in each hospital and 8 participants in each health centre in Kagera and Mwanza region were involved (Table 1).

**Table 1:** Number of Participants Required in each Region

| REGION | HEALTH FACILITY | NO OF HEALTH FACILITY | PARTICIPANT |
| --- | --- | --- | --- |
| **Mwanza** | Hospitals | 5 | 85 |
|  | H/Centre | 10 | 80 |
| **Kagera** | Hospitals | 5 | 85 |
|  | H/Centre | 10 | 80 |
| Total |  | 30 | 330 |

### Data Collection Methods/Tool

The standard Semi-structured questionnaire was developed to address socio-demographic, theoretical and practical aspects of the knowledge and practice of the participants about helping babies breathe. The questionnaire was validated and pilot tested in other hospitals and changes were made before the study.

The skilled birth attendants were assessed on knowledge, practice and demographic data of participants by using questionnaire with both closed-ended and open-ended questions. The standard questionnaires from American Heart Association were adopted and modified. The self-administered questionnaire which were composed of questions of knowledge and practice on helping babies breathe among skilled birth attendants, they also included and demographic variables like age, sex, professions, and working experiences.

The questionnaire comprised of 3 parts whereby part 1 consisted of 11 questions on demographic characteristics of the participants, part 2 consisted of 16 questions on assessing knowledge on helping babies breathe and part 3 consisted of 15 questions on assessing the practice on helping babies breathe.

### Definition of Variables and Measurements of Variable

#### Dependent Variable

**Knowledge**; is the understanding or acquiring of information through experience or education.

**Knowledge level measurement;** the subjects were asked questions on knowledge on HBB and all questions contained 16 marks. The scoring indicates that each statement consists of 5 items and correct answer carries 1 mark while the wrong answer carries zero marks. The maximum total score is 16. The knowledge level of skilled birth attendants was classified into two; adequate and inadequate knowledge. Adequate knowledge for skilled birth attendants who answered correctly at least 10 out 16 questions on knowledge, and inadequate for skilled birth attendants who correctly answered less than 10 out of 16 questions on knowledge. Standard Questionnaire was adopted and modified from American Heart Association 2005(AHA) accreditation criteria. (1).

**Practice:** it is when she/he is able to perform and follow the procedures regarding helping babies to breathe

**Practice level measurement:** The subjects were given the questions on HBB practices. The section contained 15 questions pertaining to practice and was classified into two groups that are, adequate and inadequate practice. These 15 questions were used to assess the level of practice and each correct answer was given one point. The adequate practice was for skilled birth attendants who answered correctly at least 9 out of 15 questions. With regard to inadequate practice, this involved skilled birth attendants who correctly answered less than 9 of the 15 questions(Adopted and modified from American Heart Association 2005 (AHA) accreditation criteria(15).

**Independent variable** Are those demographic factors such as age, sex, marital status, education level, experience, professional/ qualification, working facility, and training. They were measured by Descriptive measurements.

### Data Analysis Procedures

Data were analysed using SPSS version 20. Before the analysis, the cleaning of the data was done for the missing data and for checking the completeness and accuracy manually. Descriptive statistics were used to describe the demographic characteristics of the respondents and to assess the level of knowledge and practice among skilled birth attendants in rural areas of Lake zone of Tanzania. Chi-square test was used to test for association of knowledge and practice with demographic characteristics. Binary logistic regression with both odds ratios (OR) and adjusted odds ratios (AOR) was used to identify the significant factors influencing the knowledge and practice among skilled birth attendants.

### Validity and Reliability of Data

**Validity:** Randomly selected sample and employed use of a standardized questionnaire in measuring the outcome variable. Data on possible confounder was collected and adjusted during analysis.

**Reliability:** Pre-testing of the tools was done before conducting the study, to test for accuracy of the tool to yield valid information. The pilot study was carried out at Makole health center one month before the actual study. The aim of the piloting was to see how long it will take to assess skilled birth attendants and also be able to identify flows such as ambiguity in questions and establish whether or not the instructions were understandable.

### Ethical Issues

Due to the fact that the data were collected from nurses, the ethical consideration was taken into account in this study. One of the aspects of research ethics is autonomy which is about recognising the participants’ right. This was done by obtaining the informed consent of the respondents where the reasonable balance was achieved in over informing and under-informing the participants about the study exercises to be taken. It also implies that the participants exercised their autonomy to voluntarily accept or refuse to participate in the study.

The study protocol was approved by the College of Health Research Ethics Committee of the Dodoma University. Verbal and written informed consent was obtained from all participants including skilled birth attendants who were working in labour wards before the interview in voluntary basis with no compensation. The participants were informed about the purpose of the study, benefits, and risks of participating in the study and then were asked to sign. The permission to visit the health facilities was also obtained from the facility directors. The description of the study was to skilled birth attendants from the selected district hospitals and health centres in Kagera and Mwanza Region to gain access and collaboration.

## RESULTS

### Respondents Socio-Demographic Characteristics

The participants who participated in the study were all skilled birth attendants from the selected health facilities including district hospitals and health centers in each selected district. They all fell in the age category of 25-60years.

Data collected were generated from a total number of 330 respondents both males and females who participated in this study. The frequency of female respondents accounted for 218 (66.1%) while the male respondents accounted for 112 (33.9%). Based on the respondents’ age, it is observed that the highest frequency was between 25-34 years which account 167 (50.6%), followed by 35-44 years which accounted 110 respondents (33.3%), and the smallest frequency was between 45 and above which accounted for 53 respondents (16.1 Married participants formed 249 (75.5%), while 81 (24.5%) were singles (Table 2).

**Table 2:** Frequency Distribution of Socio-demographic Characteristics of Skilled Birth Attendants (N=330).

|  |  |  |
| --- | --- | --- |
| Age (years) |  |  |
| 25-34 | 167 | 50.6 |
| 35-44 | 110 | 33.3 |
| 45 and above | 53 | 16.1 |
| Sex |  |  |
| Female | 218 | 66.1 |
| Male | 112 | 33.9 |
| Marital status |  |  |
| Married | 249 | 75.5 |
| Not married | 81 | 24.5 |
| Education level |  |  |
| Certificate | 102 | 30.9 |
| Diploma | 191 | 57.9 |
| Degree | 37 | 11.2 |
| Professional/ qualification |  |  |
| Enrolled nurse | 102 | 31.0 |
| Registered nurse | 147 | 44.5 |
| AMO / MD | 81 | 24.5 |
| Work facility |  |  |
| Hospital | 170 | 51.5 |
| Health centre | 160 | 48.5 |
| Experience in labour ward |  |  |
| Less than 1 year | 100 | 30.3 |
| 1 and above years | 230 | 69.7 |
| Training on neonatal resuscitation |  |  |
| Never | 208 | 63.0 |
| Once | 73 | 22.0 |
| More than once | 49 | 15.0 |
| Equipment for neonatal resuscitation |  |  |
| Yes | 153 | 46.4 |
| No | 177 | 53.6 |

Respondents were also requested to provide their educational qualifications. The highest frequency of education was that of diploma level which accounted for191 (57.9%), followed by certificate level which accounted for 102(30.9%) and the smallest frequency is shown in the level of degree accounted for 37 (11.2%). Of 330 interviewed skilled birth attendants, registered nurses were 147 (44.5%), followed by enrolled nurses 102 (30.9%) and Doctors (AMO/MD) were 81 (24.5%) (Table 2).

From the total 330 participants out of them, 170 (51.5%) were from hospitals, and 160 (48.5%) were from health centres. Regarding the working experience of the respondents, 100 (30.3%) had less than 1year working experience, followed by 230 (69.7%) with more than 1 year working experience. Out of 330 participants, 208 (63.0%) had never attended any training/seminars on neonatal resuscitation. These were followed by 73 (22.1%) who had attended once and the smallest frequency had attended more than once 49 (14.8%). On the availability of equipment on health facilities the majority had no enough equipment 177 (53.6%) and 153 (46.4%) had enough equipment (Table 2).

### Factors Influencing Knowledge Level on Helping Babies Breathe among Skilled Birth Attendants

Chi square test and binary logistic regression were used to assess the factors associated with the knowledge level of helping babies breathe.

### Bivariate Analysis of Factors Associated with the Knowledge Level on Helping Babies Breathe among Skilled Birth Attendants

The results show that sex, education level, professional/ qualification, availability of training and working facility were significantly associated with the knowledge of helping babies breathe among skilled birth attendants (p< 0.05). The findings indicate that; for birth attendants with certificate level of education, most of them (55.9%) had adequate knowledge, while for another education level, most of the birth attendants had inadequate knowledge (61.8% and 73.0% for diploma and degree respectively). Likewise, for professional qualification, most enrolled nurses (55.9%) had adequate knowledge compared to registered nurses (40.1%) and AMO/MD (29.6%). The proportion of birth attendants with adequate knowledge was high for those working in a hospital (55.3%) compared to those working in health centres (28.8%). For both sex categories, the proportion of birth attendants with adequate knowledge was lower than the proportion of births attendants with inadequate knowledge with the proportion of adequate knowledgeable female respondents (46.3%) higher than that of male respondents (34.8%). These results are as presented in (Table 3).

**Table 3.** Cross tabulation of Knowledge Level on Helping Babies Breathe and Socio-demographic Characteristics of Skilled birth attendants (N=330).

| Demographic characteristics |  | Knowledge |  | Chi-square | P-value |
| --- | --- | --- | --- | --- | --- |
|  |  | Adequate | Inadequate |  |  |
| Age |  |  |  |  |  |
|  | 25-34 | 75 (44.9%) | 92 (55.1%) | 1.254 | .534 |
|  | 35-44 | 42 (38.2%) | 68 (61.8%) |  |  |
|  | 45 and above | 23 (43.4%) | 30 (56.6%) |  |  |
| Sex |  |  |  |  |  |
|  | Female | 101 (46.3%) | 117 (53.7%) | 4.012 | .045 |
|  | Male | 39 (34.8%) | 73 (65.2%) |  |  |
| Education level |  |  |  |  |  |
|  | Certificate | 57 (55.9%) | 45 (44.1%) | 12.537 | .002 |
|  | Diploma | 73 (38.2%) | 118 (61.8%) |  |  |
|  | Degree | 10 (27.0%) | 27 (73.0%) |  |  |
| Professional/ qualification |  |  |  |  |  |
|  | Enrolled nurse | 57 (55.9%) | 45 (44.1%) | 13.307 | .001 |
|  | Registered nurse | 59 (40.1%) | 88 (59.9%) |  |  |
|  | AMO/MD | 24 (29.6%) | 57 (70.4%) |  |  |
| Working facility |  |  |  |  |  |
|  | Hospital | 94 (55.3%) | 76 (44.7%) | 23.776 | .001 |
|  | Health center | 46 (28.8%) | 114 (71.3%) |  |  |
| Experience in labour ward |  |  |  |  |  |
|  | Less than 1 year | 40 (40.0%) | 60 (60.0%) | 0.345 | .557 |
|  | 1 and above years | 100 (43.5%) | 130 (56.5%) |  |  |
| Training on neonatal resuscitation |  |  |  |  |  |
|  | Never | 73 (35.1%) | 135(64.9%) | 14.705 | .001 |
|  | Once | 36(49.3%) | 37(50.7%) |  |  |
|  | More than once | 31(63.3%) | 18(36.7%) |  |  |
| Equipment for neonatal resuscitation |  |  |  |  |  |
|  | Yes | 94 (61.4%) | 59 (38.6%) | 1.742 | .187 |
|  | No | 96 (54.2%) | 81 (45.8%) |  |  |

### Multivariate Logistic Regression Analysis of the Factors Influencing Knowledge Level on Helping Babies Breathe among Skilled Birth Attendants

Binary logistic regression was used to identify significant predictors of knowledge among skilled birth attendants in a rural area of Lake zone of Tanzania.

The results show that working facility, professional/ qualification, and training were significant predictors of knowledge of skilled birth attendants. The odds ratios (OR) show that; birth attendants working in hospitals were 3.065 times more likely to have adequate knowledge than those working in health centers; Those who had attended training once and more than once were 1.799 and 3 respectively. Therefore, they were 185 times more likely to have adequate knowledge than those who had never attended any training; for professional/qualification. Enrolled nurses were 3 times more likely to have adequate knowledge than AMO/MD.

After adjusting for other confounders, the results show that the level of the facility, professional/ qualification, and training were significant predictors of knowledge level; the adjusted odds ratios (AOR) indicate that birth attendants working in the hospital were 3.227 times more likely to have adequate knowledge than those working in health center’s. For professional qualification, enrolled nurses were 3.118 times more likely to have adequate knowledge than other categories of birth attendants (registered and AMO/MD). For training, those who had attendant training once and more than once were each 1.778 and 3.102 times more likely to have adequate knowledge compared to those who had never attended such training.

These results are presented in (Table 4).

**Table 4:**
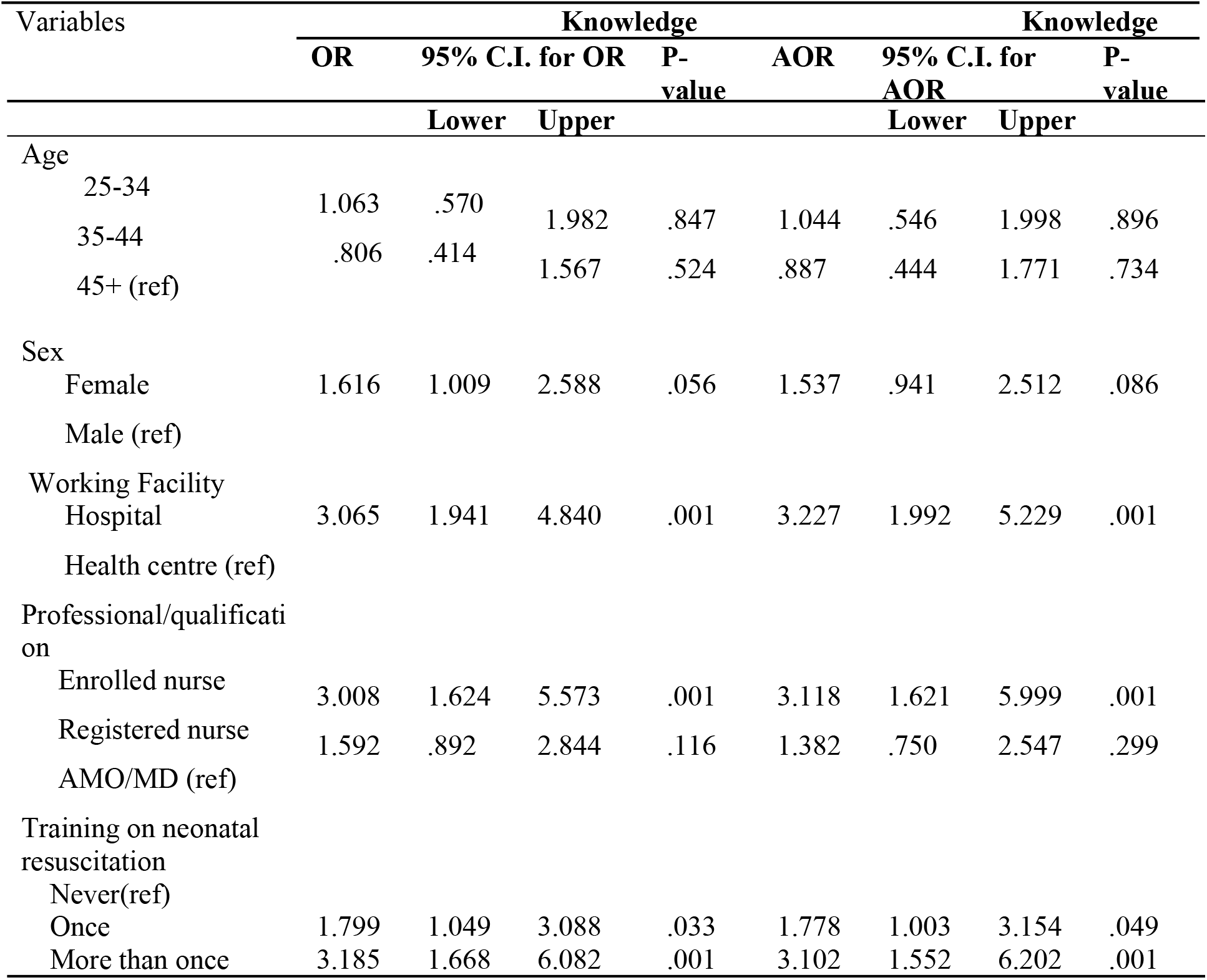
Binary Logistic Regression of Knowledge with each of the Independent Variable and all Significant Predictors of Knowledge (N=330).

### Factors Influencing Practice Level on Helping Babies Breathe among Skilled Birth Attendants

Chi square test and binary logistic regression were used to assess the factors associated with the practice level of helping babies breathe.

### Bivariate Analysis of Factors Associated with the Practice Level on Helping Babies Breathe among Skilled Birth Attendants

The results show that only working facility, experience in the labor ward and the availability of equipment, were statistically significantly associated with the practice of birth attendants among skilled birth attendants (P< 0.001). The findings further indicate that the proportion of nurses with adequate practice was high for birth attendants working in hospitals (59.4%) compared to those working in the health centers (3.8%). For experience regarding both groups, a large proportion of birth attendants had inadequate practice with those with at least one year and above of experience in labor ward (44.8%) higher than those with less than one year of experience (4.0%). For availability of equipment (49.0%) had adequate practice compared to (18.1%) with no enough equipment. The results are presented in (Table 5).

**Table 5.**
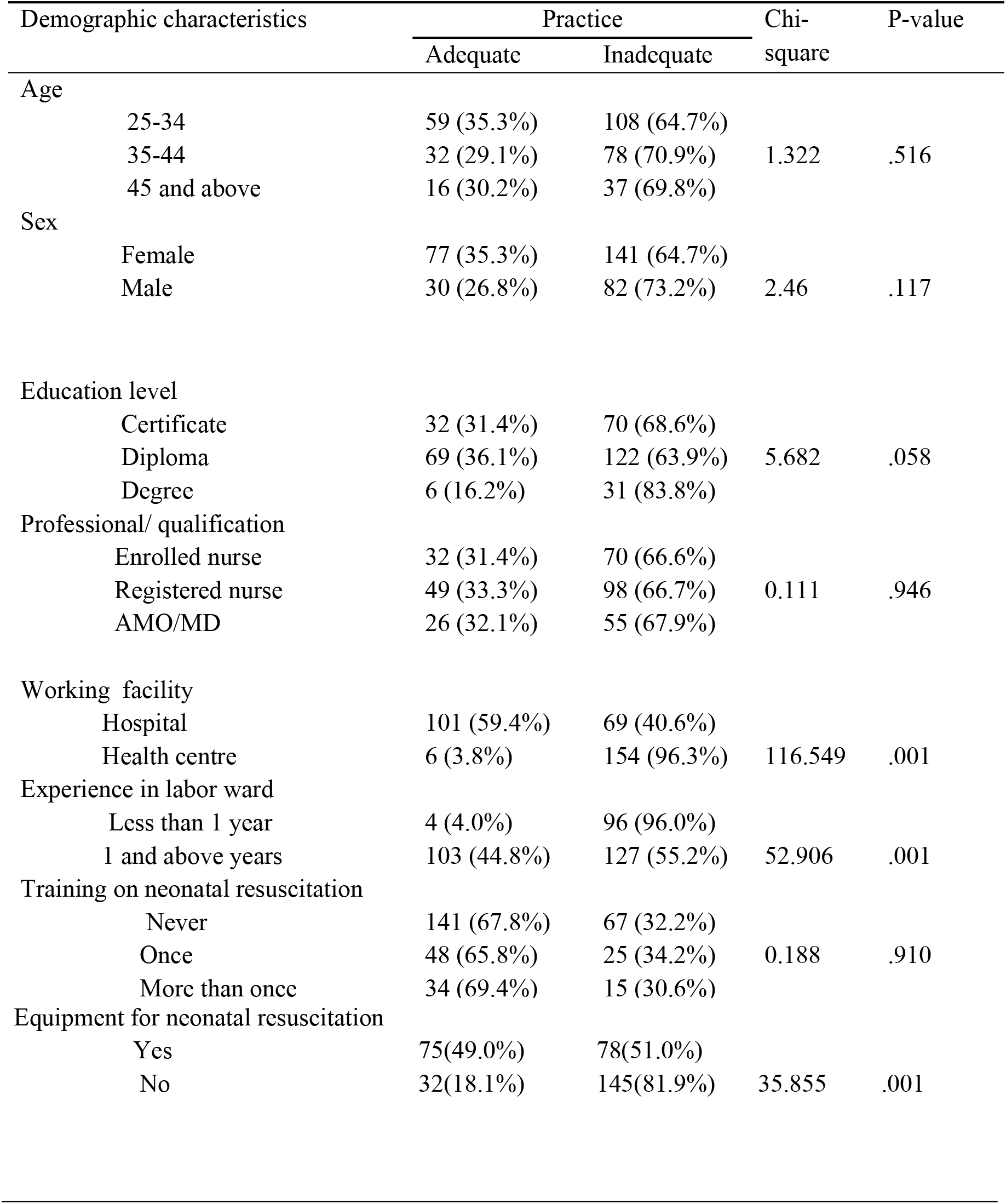
Cross tabulation Practice Level on Helping Babies Breathe and socio-demographic Characteristics of Skilled Birth Attendants (N=330)

### Multivariate Logistic Regression Analysis of Factors Influencing Practice Level on HBB among Skilled Birth Attendants

Binary logistic regression was used to identify significant predictors of practice among skilled birth attendants in a rural area of Lake Zone of Tanzania. The results show that the level of facility, experience, and availability of equipment were predictors for practice.

The odds ratios (OR) show that birth attendants working in the hospital were 37.57 times more likely to have adequate practice than those working in health centres. Birth attendants with at least one year of experience were 19.465 times more likely to have adequate practice than those with less than one year of experience in the labour ward. Those with enough equipment were 4.3567 times more likely to have adequate practice than those with not enough equipment.

After adjusting the other factors, the results show that the level of the facility, experience in the labour ward and enough equipment were significant predictors of practice. The adjusted odds ratios indicate that birth attendants working in the hospital were 43.807 times more likely to have adequate practice than those working in health centres. For experience, birth attendants with at least one year of experience were 15.418 times more likely to have adequate practice than those with less than one year of experience. For enough equipment, those who had enough equipment were 4.355 times more likely to have adequate practice than those working in facilities with no enough equipment. The results presented in table 6.

**Table 6.**
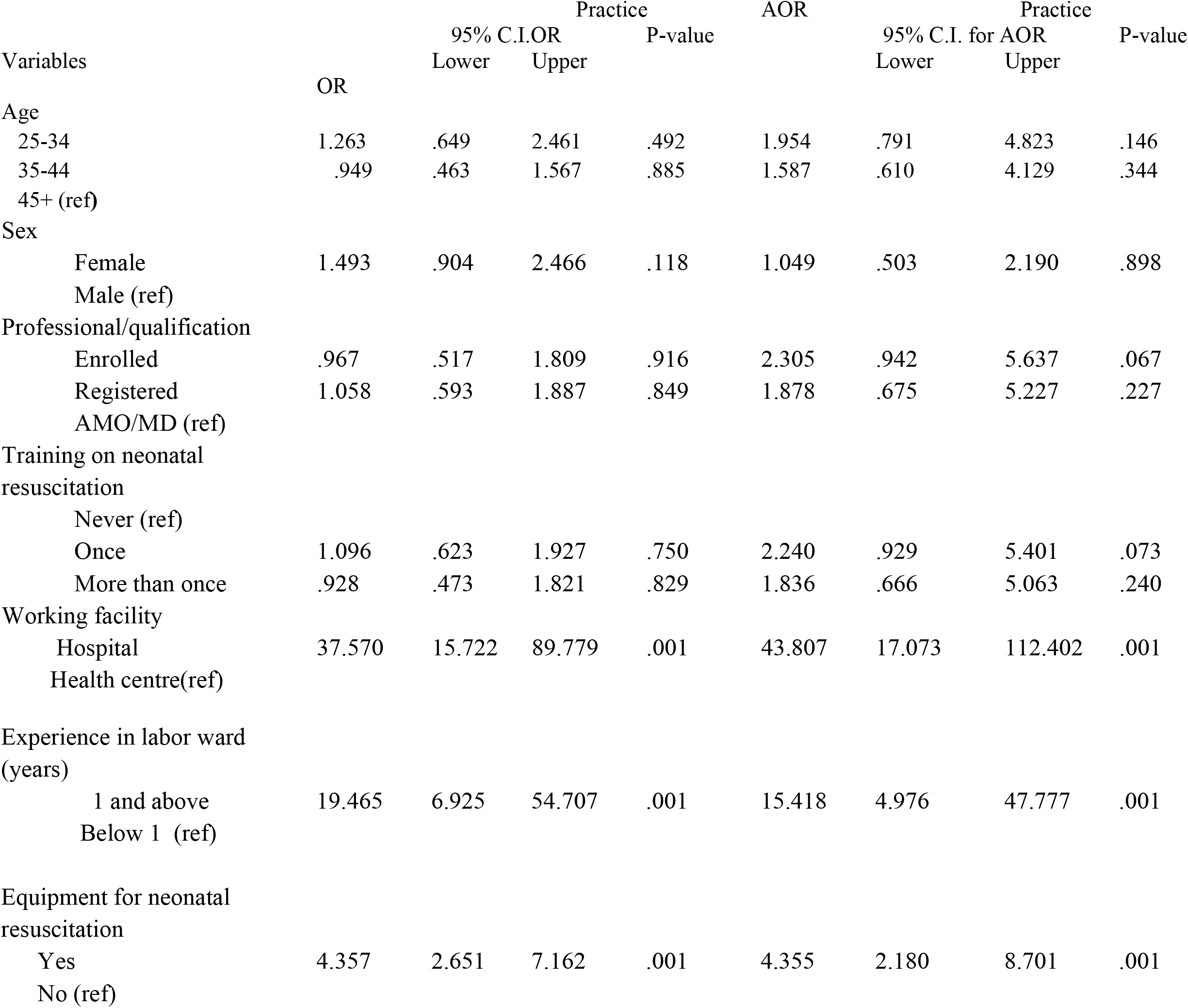
Binary Logistic Regression Practice with each of Independent Variable and all Significant Predictors of Practice (N=330).

## DISCUSION

This chapter discusses Factors Influencing Knowledge Level and practices on Helping Babies Breathe among Skilled Birth Attendants. The findings are based on a representative sample of the skilled birth attendants and adopted standard measures of factors influencing knowledge and practice level on helping babies breathe among skilled birth attendants.

### Factors Influencing Knowledge Level on Helping Babies Breathe among Skilled Birth Attendants

The results show that, working facility, professional/ qualification and training on neonatal resuscitation were statistically significantly associated with the knowledge of helping babies breathe among skilled birth attendants (p< 0.05).

The factor which was observed to contribute to this problem was a shortage of staff working in the labour ward, low frequency of neonatal training, lack of refresher training on helping babies breathe. This could be due to lack of skilled birth attendants working in labor ward and majority did not attend training on neonatal resuscitation. So the regular training and increasing number of skilled birth attendants in labor ward was needed.

The current study was similar with other findings by (16) show that acquisition of knowledge and skills was associated with the level of training and the facility type in which birth attendants were working. They also showed that birth attendants from lower cadres (e.g., midwives) improved significantly more from pre-training to post-training than those from higher cadres (e.g., physicians). This could be due to majority of skilled birth attendants were working in labor ward and those who attending training on neonatal resuscitation have adequate knowledge on neonatal resuscitation as compared to physician.

The findings of this study are linked with the study done in Afghanistan by (17) whose results showed that; facility type, infrastructure, years of experience offering, training on new-born resuscitation, and confidence in performing new-born resuscitation had a statistically significant association with knowledge of health professionals on neonatal resuscitation. Trained health professionals (respondents) had more than 26 times sufficient knowledge on neonatal resuscitation than untrained health professionals or untrained health professionals. This might be due to those who attending training on neonatal resuscitation had sufficient knowledge also those with more experience they practice more on neonatal resuscitation as compare to those with less experience in labor ward.

Our findings are consistent with (18) who also found that lack of neonatal resuscitation training was among factors that influence knowledge. This might be due to shortage of staff.

### Factors Influencing Practice Level on Helping Babies Breathe among Skilled Birth Attendants

The findings of the current study show that working facility, experience in the labour ward and availability of equipment was statistically significantly associated with the practice level on helping babies breathe among skilled birth attendants. The factor which was observed to contribute to this problem was a low number of staff working in the labour ward, inappropriate space for neonatal resuscitation, not enough time to practice due to many activities. This could be due to failure to follow guideline for neonatal resuscitation and lack of adequate knowledge on neonatal resuscitation.

Furthermore, the current study was similar with a study done by(19) shown that lack of equipment is common the reason for not being able to practice helping babies breathe. These observed that those with enough equipment have appropriate practice as compared to those with no enough equipments.

These findings are consistent with a study done in Ethiopia which reported that training and well equipped health faculty had a statistically significant association with the practice of health professionals on neonatal resuscitation. Untrained health professionals were 7 times with less practice level than trained respondents (health professionals) on neonatal resuscitation and well equipped health facility had a statistically significant association with the practice of health professionals on neonatal resuscitation well equipped health facility were 4 times better practice level than unwell equipped health facility on neonatal resuscitation(20).

## CONCLUSSION

Regarding the findings of the current study, it was revealed that working facility, Professional/ qualification, and training was a significant predictor of knowledge while working facility, experience and equipment’s was significant predictor of practice. There is a need to find effective measures on how to reduce those factors, which affect knowledge and practice on helping babies breathe.

## Acknowledgement

We would like to acknowledge the support given by the Ministry of Health in Tanzania for their sponsorship, and my deeply sincere thanks to my supervisors Dr. Secilia Ng’weshemi Kapalata and Dr. Alex Ernest, for their tireless efforts, patience, professional guidance, and advice. Lastly, I would like to thank all those who responded to the questionnaire.

## Author contributions

**Conceptualization**: Cecilia B.Mzurikwao, Secilia Kapalata Ng’weshemi, Alex Ibolinga Ernest

**Data collection**: Cecilia B.Mzurikwao.

**Formal analysis:** Cecilia B.Mzurikwao,Secilia Kapalata Ng’weshemi,Alex Ibolinga Ernest.

**Investigation:**Cecilia B.Mzurikwao

**Methodology:** Cecilia B.Mzurikwao, Secilia Kapalata Ng’weshemi,Alex Ibolinga Ernest

**Supervision**: Secilia Kapalata Ng’weshemi,Alex Ibolinga Ernest

**Writing –original draft**: Cecilia B.Mzurikwao, Secilia Kapalata Ng’weshemi,Alex Ibolinga Ernest

**Writing –review & editing**: Cecilia B.Mzurikwao,Secilia Kapalata Ng’weshemi,Alex Ernest

